# Mitochondria as Indispensable Yet Replaceable Components of Germ Plasm: Insights into Primordial Germ Cell Specification in Non-Teleost Sturgeons

**DOI:** 10.1101/2025.01.05.631362

**Authors:** Linan Gao, Roman Franěk, Tomáš Tichopád, Marek Rodina, David Gela, Radek Šindelka, Taiju Saito, Martin Pšenička

**Affiliations:** Faculty of Fisheries and Protection of Waters, South Bohemian Research Center of Aquaculture and Biodiversity of Hydrocenoses, University of South Bohemia in Ceske Budejovice, Zatisi 728/II, 389 25 Vodnany, Czech Republic; Laboratory of Non-Mendelian Evolution, Institute of Animal Physiology and Genetics of the CAS, Liběchov, 277 21, Czech Republic; Laboratory of Gene Expression, Institute of Biotechnology, BIOCEV, Vestec, Czech Republic; Nishiura Station, South Ehime Fisheries Research Center, Ehime University, 25-1 Uchidomari, Ainan-cho, Ehime 798-4206, Japan

**Keywords:** Primordial germ cells, Mitochondria, Matrilineal genetics, Germ plasm, Microinjection, Sturgeon reproduction

## Abstract

While it is widely recognized that mitochondria are components of germ plasm, their specific role in the formation and specification of primordial germ cells (PGCs) remains poorly understood. Furthermore, it has not been established whether mitochondria in germ plasm possess unique characteristics essential for their function. In this study, we demonstrate that mitochondria are indispensable for PGCs development in non-teleost fishes and that their role is not dependent on their origin from germ plasm. Using sturgeon embryos, we showed that UV radiation applied to the vegetal pole effectively eliminates germ plasm, including mitochondria, and prevents PGCs formation. Remarkably, we restored germ plasm function and PGCs development by injecting mitochondria derived from donor eggs, even when these mitochondria were not originally part of the germ plasm. Transplanted mitochondria were successfully identified in larval PGCs using a fluorescent PKH26 tracer, and in interspecies transplantation experiments, their presence was confirmed using species-specific mtDNA and mtRNA primers in larvae and individual PGCs. Our findings reveal that mitochondria are critical but not germ plasm-specific determinants of PGCs formation. This study provides novel insights into the developmental pathways of germ cells and establishes a previously unrecognized flexibility in mitochondrial functionality within the germline. These findings also offer a potential method for conserving matrilineal genetics in critically endangered species like sturgeons, while simultaneously opening new avenues for studying germlines with high interspecies mitochondrial heteroplasmy and contributing to broader evolutionary and conservation biology.

## Introduction

Sturgeon, belonging to the order Acipenseriformes, represent a lineage of fishes whose origins can be traced back to the Lower Jurassic period, highlighting their profound evolutionary heritage (Bemis et al., 1997). In recent centuries, however, their natural populations have faced severe declines. Those declines are attributed to a confluence of factors including overexploitation through poaching for their prized caviar, the adverse effects of water pollution, habitat degradation, and notably, the construction of dams that obstruct their reproductive migration (Vick & Kelly, 2021). The International Union for Conservation of Nature has identified sturgeons as one of the most endangered group of animals on earth (Nicki et al., 2010). Given this critical status, the urgency to conserve their dwindling diversity and foster the recovery of these ancient, majestic fishes cannot be overstated.

Cryopreservation offers a method for the storage of living cells for a virtually infinite period. The cryopreservation of sperm, eggs and embryos in a wide variety of species is well documented and these techniques have found utilization in breeding and conservation programmes (Martínez-Páramo et al., 2017). While sperm has been successfully cryopreserved in several fishes including sturgeons (Kopeika et al., 2007; Kumari & Maurye, 2021; Tiersch et al., 2007), we still lack efficient and high throughput techniques for cryopreserving fish eggs and embryos. Challenges for this include the large size of the oocyte, high yolk content, and limited permeability to cryoprotectants (Mazur et al., 2008). Moreover, the failure to achieve successful surrogate reproduction in sturgeon—a promising conservation strategy—further exacerbates the challenges of safeguarding maternal genetic material. These technological bottlenecks place the maternal genetics, which generally include maternal mitochondria, at high risk. Addressing this, there is an urgent need to establish a new technology to conserve sturgeon maternal genetics.

The primordial germ cells (PGCs) of the sturgeon are exclusively determined and formed from the maternal germ plasm (Saito et al., 2014). Mitochondria, as a key component of the germ plasm (Beams & Kessel, 1974; Eddy, 1975), are abundant and directly involved in germ line formation (Amikura et al., 2001; Kalt, 1973; Saffman & Lasko, 1999). Despite the recognition of mitochondria as a critical constituent of germ plasm, their precise role in the formation and specialization of PGCs remains unclear (Reunov et al., 2019). In addition, there are no studies exploring whether mitochondria function independently of the overall role of the germ plasm, nor their specific contribution to supporting the development of PGCs. (Saito et al., 2008) (Yoshizaki & Lee, 2018) (de Siqueira-Silva et al., 2021; Franěk et al., 2021; Hattori et al., 2019; Kawamura et al., 2024; Okutsu et al., 2007; Yutaka Takeuchi et al., 2004)(PšeniČka et al., 2015)(Baloch et al., 2019; Linhartová et al., 2015)(Saito et al., 2018)(Romney et al., 2023)In this study, we took advantage of the unique developmental pathway of PGCs in sturgeon which we identified previously. The PGCs are exclusively formed from maternal germ plasm located in the vegetal pole of the embryo, while the rest of the vegetal hemisphere is utilized exclusively as endogenous nutrition in the form of yolky cells (Saito et al., 2014; Shah et al., 2022, 2024). Irradiation of the vegetal pole using UV light leads to germ plasm depletion and a loss of PGCs, without compromising the viability of the embryos (Saito et al., 2018). This experimental model provides a unique opportunity to investigate the specific roles of mitochondria in germ plasm function and PGC formation.

Our study has developed and applied an innovative technique—interspecies replacement of mitochondria in sturgeon germ cells. This involved eliminating germ plasm including mitochondria in the vegetal pole of the host embryo using UV irradiation, isolating mitochondria from donor sturgeon eggs, and transplanting these mitochondria into the vegetal pole of host embryos (Fig. 1). This approach aims to elucidate the specific roles of mitochondria in germ cell development. Notably, our findings demonstrate for the first time that mitochondria do not require origin-specific properties to support PGC development, challenging prior assumptions about germ plasm-specific mitochondrial specialization. Moreover, the introduction of mitochondrial replacement in germ plasm enables the incorporation of maternal genetics into conservation strategies, which is not possible through other means, and provides new tool for the conservation and reproduction of endangered species such as sturgeon. Additionally, it also opens new avenues for research on germlines with high interspecies mitochondrial heteroplasmy, benefiting fields such as evolutionary biology, genetics and genomics, developmental biology and conservation biology.

**Fig. 1.**
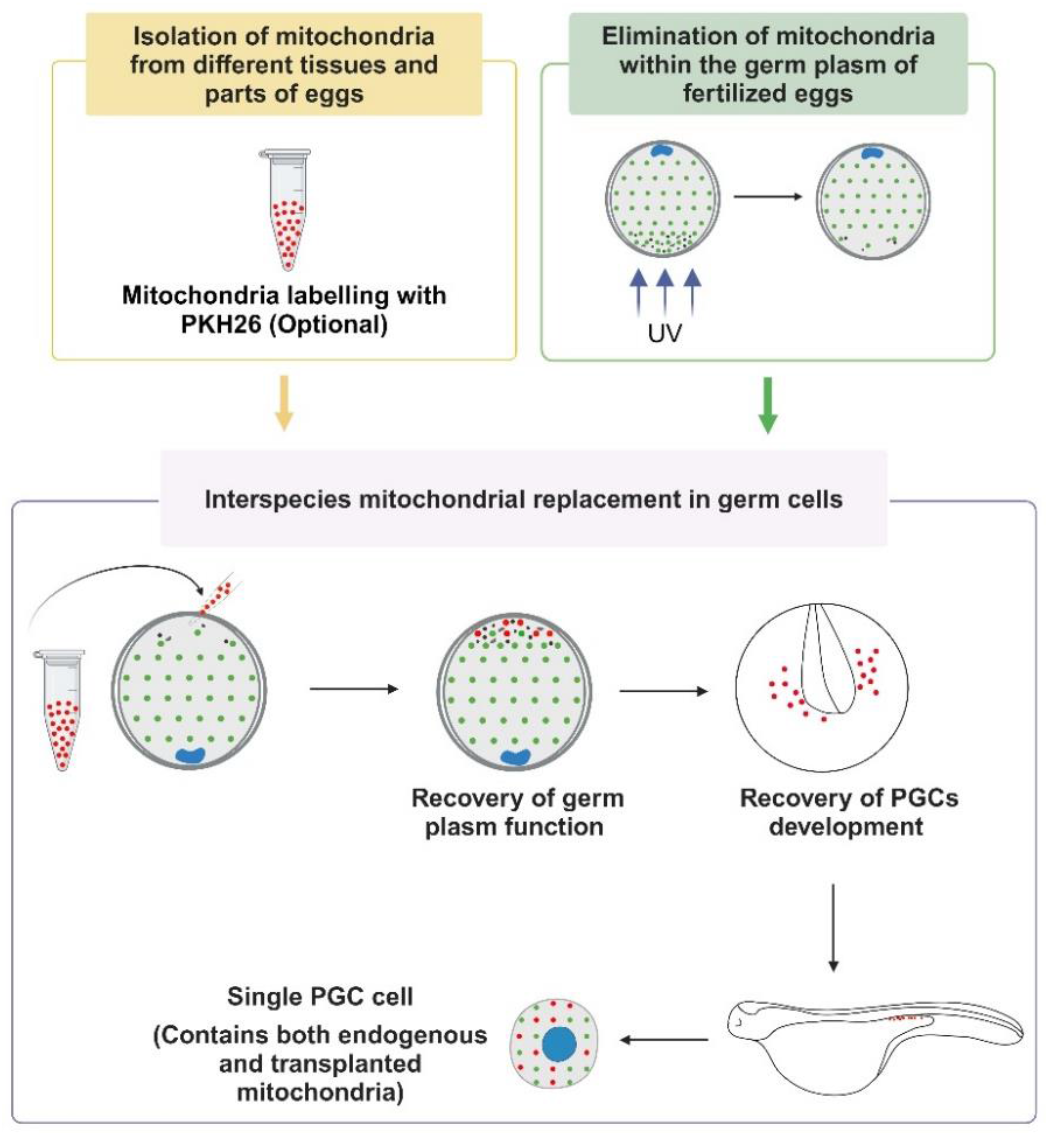
Schematic of study design. The blue colour represents the nucleus. The red and green colours represent the donor and the host mitochondria. The areas with different black shapes are the germ plasm. The figure was created with BioRender.

## Materials and methods

### Ethics

The experiments took place at the Faculty of Fisheries and Protection of Waters, Research Institute of Fish Culture and Hydrobiology, University of South Bohemia in České Budějovice, Czech Republic. All experimental procedures were performed in accordance with national and institutional guidelines on animal experimentation and care. This study was approved by the Animal Research Committee of the University of South Bohemia in České Budějovice. Fish were maintained according to the principles of animal welfare and principles of laboratory animal care based on the Guidelines for the Use of Animals in Research.

### Isolation and Purification of Mitochondria

Crude mitochondria were isolated by double centrifugation using a Mitochondria Isolation Kit (Cat# MITOISO1, SIGMA-ALDRICH^®^) according to the manufacture’s protocol. Briefly, fresh egg, liver and muscle samples, as well as vegetal and animal poles obtained from whole eggs using microcapillary, were homogenized with extraction buffer containing 2 mg/ml albumin (Cat# A0474, SIGMA-ALDRICH^®^). The homogenate was centrifuged at 600 × g for 5 min at 4 °C. The resulting supernatant was removed and centrifuged at 11,000 × g for 10 min at 4 °C. The supernatant was discarded, and the pellet was resuspended in an extraction buffer. This two-step differential centrifugation process was repeated. The final mitochondrial pellet was resuspended in a storage buffer (50 mM HEPES, pH 7.5, containing 1.25 M sucrose, 5 mM ATP, 0.4 mM ADP, 25 mM sodium succinate, 10 mM K_2_HPO_4_, and 5 mM DTT) at 40 mL per 100 mg tissue. The protein concentration was determined using the BCA protein assay kit (Cat# 23225, ThermoFisher SCIENTIFIC) and adjusted to 2-3 mg/ml.

The crude mitochondria suspension was further purified by Percoll density gradient centrifugation according to previously established protocol (Nukala et al., 2006) with modifications. Stocks of 40 % and 24 % Percoll (Cat# P1644, SIGMA-ALDRICH^®^) were prepared fresh in a storage buffer. An equal volume of 24 % Percoll was added to the mitochondria suspension to get a 12 % final Percoll concentration. A volume of 3.5 ml of 40 % Percoll, 24 % Percoll and 12 % Percoll with mitochondria were carefully loaded in 12 ml ultra-clear tubes in sequence. The tubes were then centrifuged in a swinging bucket rotor type SW 32 Ti at 30,700 × g for 10 min at 4 °C in an Optima L-90K ultracentrifuge (Beckman Coulter). A yellowish turbid band that formed at the interphase of the 40 % and 24 % layers was carefully collected and placed into fresh tubes. The tube was topped off with storage buffer and centrifuged at 16,700 × *g* for 10 min at 4 °C. The supernatant was removed, and the pellet was resuspended in a fresh tube with storage buffer. The tube was centrifuged at 11,000 × *g* for 10 min at 4 °C. The final mitochondrial pellet was resuspended to approximately 1 mg/ml in storage buffer.

### Labelling of mitochondria

Mitochondria were labelled by using a PKH26 Red Fluorescent Cell Linker Kit for General Cell Membrane Labelling (Cat# MINI26, SIGMA-ALDRICH^®^) according to the manufacture’s protocol with modifications. Briefly, a 2 × Dye Solution (8 × 10^−6^ M) in storage buffer was prepared. An equal volume of mitochondrial suspension was rapidly added to the 2 × Dye Solution and immediately mixed by pipetting. The mitochondria/dye suspension was incubated for 5 min with periodic mixing. Then 1% bovine serum albumin (BSA; Cat# A8806, SIGMA-ALDRICH^®^) was added and incubated for 1 min to stop the staining by binding of excess dye. The suspension was centrifuged at 11,000 × *g* for 10 min at 4 °C and the supernatant was carefully removed. The pellet was resuspended in storage buffer and centrifuged at 11,000 × *g* for 10 min at 4 °C to remove unbound dye. After the final wash, the pellet was resuspended in storage buffer for later injection.

### Cryopreservation of mitochondria

The protocol used for cryopreservation of mitochondria was according to Nukala et al. (Nukala et al., 2006) with modifications. The suspension of mitochondria in storage buffer was mixed with dimethyl sulfoxide (DMSO, 10 %, v/v, Cat# 472301, SIGMA-ALDRICH^®^) in 1.8 ml cryotubes. Then the tubes were immediately placed into a -80 °C freezer. For further use, the cryotube was thawed in a water bath at 37 °C. When the sample was completely thawed, the tube was centrifuged at 600 × *g* for 5 min at 4 °C. Then the supernatant was carefully transferred and centrifuged at 11,000 × *g* for 10 min at 4 °C to remove the cryomedium. The supernatant was discarded, and the pellet was resuspended in storage buffer, washed once more and finally stored in a storage buffer at 4 °C until use.

### Preparation of Host Embryos

Gametes were collected from sterlet according to Saito and Psenicka (Saito & Psenicka, 2015). After fertilization, the embryos were treated with 0.1 % tannic acid three times for 2 min each to remove their stickiness, then washed with filtered water three times. The outer chorion layer was removed using fine forceps under a stereomicroscope. The embryos were placed on a UV transilluminator (TFS 26V, UVP, Analytik Jena), and their vegetal hemispheres were irradiated with 254nm wavelengths for 2min with total dose of 360 mJ/cm^2^ according to Saito et al. (Saito et al., 2018). To confirm germ plasm destruction, PGCs were labelled by injecting the vegetal pole with 2% FITC-dextran (Cat# FD500S, SIGMA-ALDRICH^®^, MW = 500,000) dissolved in 0.2 M potassium chloride (Cat# 16200-31000, PENTA). FITC-dextran was used based on the findings of Saito et al. (Saito & Psenicka, 2015), who demonstrated that injecting FITC-dextran into the vegetal pole specifically labels PGCs. The number of PGCs was assessed using a fluorescent microscope at the late neurula stage in both irradiated and controlled embryos. Several embryos were also fixed at the 256-cell stage and examined by electron microscopy (see below).

### Transplantation of mitochondria into germ plasm of embryos

The embryos were placed on an agar-coated (1 %) dish filled with dechlorinated tap water. Mitochondria isolated from eggs and tissues of sterlet, and Siberian sturgeon were mixed with 2 % FITC-dextran (as explained above) and microinjected into the vegetal pole of 1 - to 4 - cell stage UV-irradiated sterlet embryo. In order to track the injected mitochondria, they were labelled by PKH26 prior to microinjection. Then the embryos were incubated in an incubator at 16 °C in dechlorinated tap water with two water changes per day. The number of PGCs was assessed under a fluorescent microscope at the late neurula stage (note: each group consisted of two to three females, with a minimum of 100 embryos per female; each embryo was injected with approximately 5 nl of mitochondria per injection).

### Histology and Microscopy

We used transmission electron microscopy to observe and compare the composition of purified and unpurified mitochondrial suspensions, as well as the ultrastructure of the vegetal pole germplasm in the intact embryos (control) and the UV-irradiated and mitochondria-injected groups. Mitochondrial suspensions and embryos at the 256-cell stage were fixed with 2.5% glutaraldehyde (Cat# G5882, SIGMA-ALDRICH^®^) in phosphate buffered saline (Cat# P4417, SIGMA-ALDRICH^®^) for 2 h and dehydrated through an acetone series, and finally embedded in Poly/Bed^®^ 812 Embedding Media (Cat# 08791-500, Polysciences). Then a series of ultrathin sections of mitochondrial suspension and embryos at the vegetal pole along the animal-vegetal axis were cut using a Leica UCT ultramicrotome, double stained with uranyl acetate and lead citrate, mounted on a grid, and observed using a transmission electron microscopy (TEM, JEOL 1010, JEOL Ltd).

For observation of the PGCs with transplanted mitochondria, 5 days post hatch (dph) larvae, corresponding to 12 days post fertilization (dpf), were prepared for plastic sections according to Sullivan-Brown et al. (Sullivan-Brown et al., 2011). Briefly, samples were fixed with 4 % Paraformaldehyde (Cat# 23700-31000, PENTA) for 24 h at 4 °C and then dehydrated in an ethanol series and embedded in JB-4 resin (Cat# EM0100, JB4 embedding kit). Specimens were sectioned to 8 μm using a rotary microtome (Leica Biosystems). Sections were stained with Fluoroshield with DAPI (Cat# F6057, SIGMA-ALDRICH^®^) and observed and imaged using a fluorescent microscope (BX51, Olympus).

### Identification of transplanted mitochondria

DNA was extracted from 10dph sterlet larvae, and from 10dph sterlet larvae injected with mitochondria of Siberian sturgeons using a AllPrep DNA/RNA Mini Kit (Cat# 80004, Qiagen). PCR amplification was then performed using mtDNA specific primers (Mugue et al., 2008). The amplification conditions were 95 °C for 2 min; 35 cycles of 92 °C for 20 s, 57 °C for 30 s, and 72 °C for 30 s, and last synthesis at 72 °C for 10 min. The PCR product was checked for positive amplification on 2 % agarose gel.

### Design and validation of species-specific qPCR

Universal primers (5’ - TCCAACATCTCCGTGTGATGAA - 3’ and 5’ - TGTTGGGTTGTTTGATCCTGTTTG - 3’) for sterlet species were designed for cytochrome B using the NCBI database (NC_027417.1:14420-15560, OK094493.1 and GU647227.1). Total RNA was extracted from sterlet and Siberian sturgeon eggs using the RNeasy Mini Kit (Cat# 74104, Qiagen). Extracted RNA was processed for cDNA synthesis using WizScript™ RT FDmix (Cat# W2205, Hexamer), thermal cycler programmed at 25 °C for 10 min, 42 °C for 30 min, 85 °C for 5 min, and held at 4 °C. The amplification conditions were 94 °C for 1 min and 35 cycles of 94 °C for 15 s, 59 °C for 30 s, and 72 °C for 30 s with a final extension of 72 °C for 10 min. The amplified product was checked for positive amplification on 2 % agarose gel. The amplicon was cut with a sterile scalpel from the gel and purified using the GeneJET Gel Extraction Kit (Cat# K0691, ThermoFisher SCIENTIFIC), followed by Sanger’s sequencing. Using Sanger Products, pair-wise SNP were identified and used for probes design (Supplementary Note). We designed primers and TaqMan probes that attach to a conserved mtDNA region using DNAMAN and Primer Premier software, thus equally amplifying both mitotypes.

Mitochondrial DNA was isolated using a Qiagen kit from collected pure species-specific mitochondria. dPCR was prepared using 4 µl of 50 or 5 pg of mitochondrial DNA, 1 µl of probe (final concentration 0.25 µM), 1.8 µl of primer mix (450 nM), 10 µl of ddPCR Supermix for probes (Cat# 1863024, Bio-Rad) and 2.2 µl of RNase-free distilled water (Cat# 10977015, ThermoFisher). Temperature protocol for dPCR was: predenaturation at 95 °C for 10 min, followed by 40 cycles of 94 °C for 30 s and 58 °C for 60 s. The last step was heating for 10 min at 98 °C and cooling to 4 °C. Droplet analysis was performed using ABS setting.

### Single PGCs analysis

The PGCs visualized by FITC-dextran and PKH26 were isolated manually from sterlet embryos injected with the mitochondria of Siberian sturgeon at the late neurula stage and photographed using a fluorescent microscope (IX83, Olympus). The cells were then stored in nuclease-free water with 1 % BSA at -80 °C for identification. Single PGC were collected in 5 μl of RNase-free distilled water. dPCR was prepared using a complete volume of lysed PGC, 1 µl of probe (final concentration 0.25 µM), 1.8 µl of primer mix (450 nM), 10 µl of ddPCR Supermix for probes and 2.2 µl of RNase-free distilled water. Temperature protocol for dPCR was predenaturation at 95 °C for 10 min, followed by 40 cycles of 94 °C for 30 s and 58 °C for 60 s. The last step was heating for 10 min at 98 °C and cooling to 4 °C. Droplet analysis was performed using ABS setting.

### Statistical analysis

Data on mitochondrial number in germ plasm and PGCs number in embryos were compared between the groups using Kruskal-Wallis test followed by Dunn’s multiple comparisons. *P*-values < 0.05 was considered statistically significant.

## Results

### 1. Ultraviolet irradiation eliminates germ plasm

We have confirmed that UV irradiation of the vegetal pole leads to the elimination of the germ plasm (Fig. 2A and Fig. 2B) and a subsequent decline in the numbers of PGCs. The impact of UV irradiation on mitochondria was analysed by counting the number of mitochondria in the germ plasm in each experimental group. A significant (*p* < 0.05) decrease in mitochondrial numbers was observed in the UV-irradiated group (Fig. 2A, right). The mitochondria in the germ plasm of nearly all embryos were destroyed and only a few mitochondria remained visible (Fig. 2A, UV). However, the number of mitochondria in the germ plasm increased significantly (*p* < 0.05) after injecting egg mitochondria into UV-irradiated embryos (Fig. 2A, right). Interestingly, mitochondrial injection can recover not only the mitochondria themselves, but the entire functionality of germ plasm including PGCs migration.

**Fig. 2.**
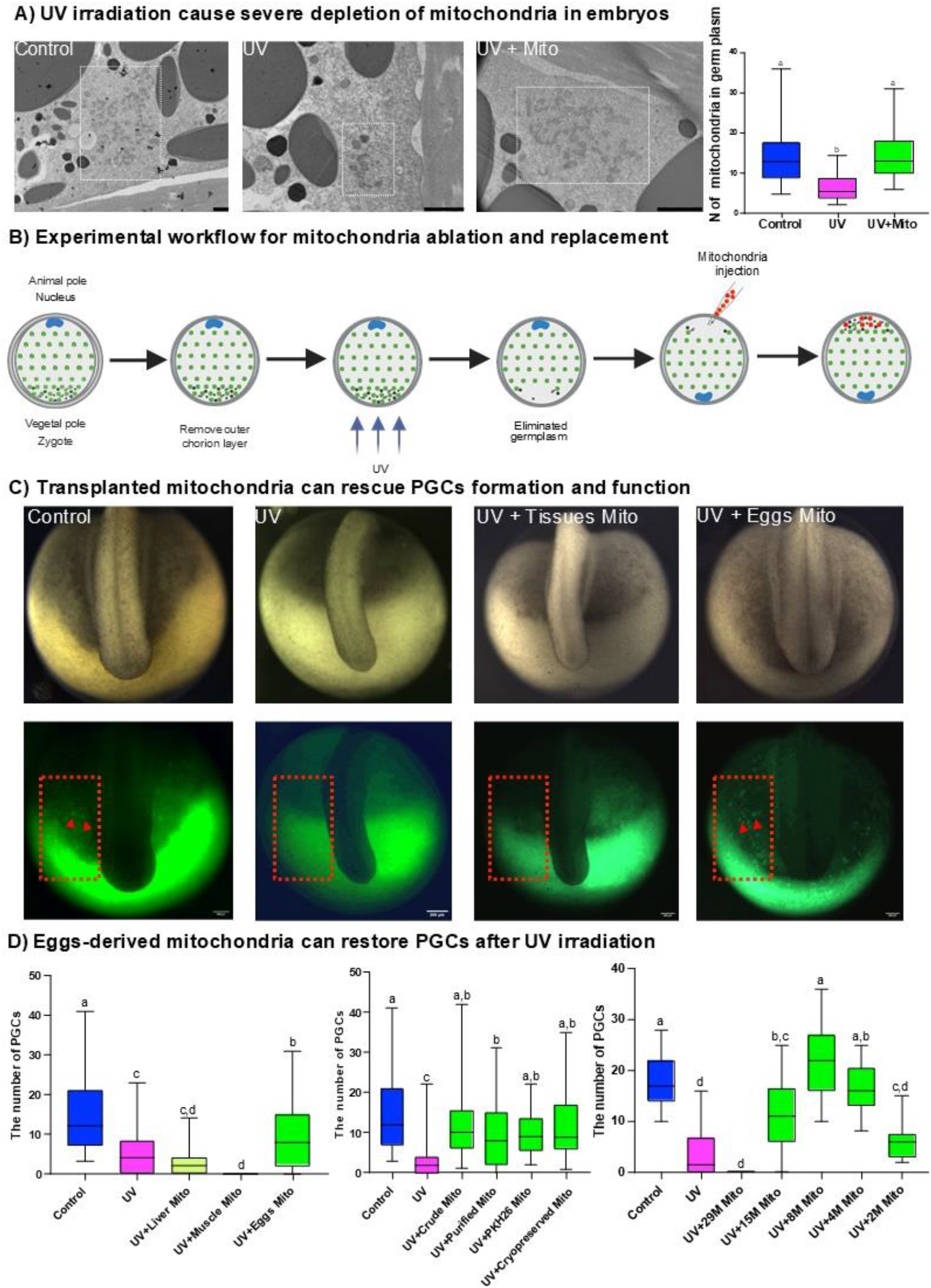
Only egg-derived mitochondria can successfully rescue PGCs depleted with UV irradiation. (A) Left: TEM image of germ plasm with mitochondria (boxed areas) in vegetal pole germ plasm along the animal-vegetal axis at the 256-cell stage. Control: The germ plasm in a control embryo, which contains many mitochondria. UV: The germ plasm in a UV-irradiated embryo, which showed a decrease of mitochondria when compared to the control. UV+Mito: The germ plasm in a UV-irradiated and then egg mitochondria-injected embryo, which showed a recovery of mitochondria when compared to the UV group. Right: Comparison of the mitochondrial number in germ plasm in different treatment groups, *n* = 40, 43 and 82 sample points. (C) Control: Control embryo. UV: UV-irradiated embryo. UV+Eggs Mito: UV-irradiated and eggs mitochondria-injected embryo. UV+Tissues Mito: UV-irradiated and tissues mitochondria-injected embryo. Box plots: area with FITC-dextran labelled PGCs. The arrow points to some of the FITC-labeled PGCs. (D) Left: Comparison of PGCs numbers in different tissues purified mitochondrial injection groups, *n* = 109, 70, 50, 42 and 92 embryos. Middle: Comparison of PGCs numbers in different eggs mitochondrial treatment groups, *n* = 109, 137, 86, 92, 41 and 43 embryos. Right: Comparison of PGCs numbers in eggs mitochondrial injection groups with different quantities. Approximately 29, 15, 8, 4 and 2 ×106 mitochondria were injected, respectively. *N* = 41, 40, 14, 40, 41, 40 and 41 embryos. Cross lines in all box plots represent median values, and different lowercase letters indicate significant differences (*P* < 0.05 by Kruskal-Wallis test followed by Dunn’s multiple comparisons). Scale bar in (A) indicates 2 μm, and 200 μm in (C).

### 2. The effectiveness of mitochondria in rescuing primordial germ cells

Crude mitochondria were obtained by a simple double centrifugation method. This method yields a mitochondrial suspension contaminated with other cellular components (Supplementary Fig. 1). The crude mitochondria from eggs (whole egg, vegetal pole, and animal pole) and tissues (liver and muscle) were separately injected into the vegetal pole of UV-irradiated embryos to compare their ability to recover the germ plasm and subsequent PGCs development. The UV-irradiated embryos and embryos injected with mitochondria derived from liver and muscle tissue had no or very few (Average 4) PGCs (Fig. 2C). In contrast, embryos injected with mitochondria derived from oocytes contained a comparable number (Average 14) of PGCs to control embryos (Fig. 2C). Furthermore, we found that the injection of mitochondria extracted from different parts of the egg into UV-irradiated embryos did not make a difference in the recovery of PGCs. On average, 14, 13, and 13 PGCs were observed in embryos injected with mitochondria from whole eggs, the vegetal pole, and the animal pole, respectively. Therefore, in subsequent experiments we used mitochondria extracted from the whole egg.

To exclude the presence of other possible factors in the injected suspension of mitochondria responsible for the recovery of PGCs, we introduced a purification step of mitochondria by a density gradient centrifugation. This approach resulted in markedly purer mitochondria suspension, predominantly consisting of mitochondria (Supplementary Fig. 1). Subsequently, we repeated the injection with purified eggs and tissues (liver and muscle) mitochondria into the UV-irradiated embryos to compare the recovery of PGCs by counting their number. A very similar result was obtained, showing that only injection of mitochondria from eggs could rescue germ plasm (Fig. 2D, left). Compared to the control group, the UV-irradiated group exhibited a significant (*p* < 0.05) reduction in the number of PGCs and the number of PGCs did not increase after liver and muscle mitochondrial injection. However, PGCs recovery was observed after the injection of eggs-derived mitochondria.

Then, to verify whether different treatments of egg mitochondria affect the recovery of germ plasm, we compared injection of freshly prepared crude mitochondria, purified mitochondria, PKH26 labelled mitochondria and cryopreserved mitochondria into the UV-irradiated embryos. The recovery of PGCs was observed in all injection groups (Fig. 2D, middle). This implies that labelling with PKH26 and cryopreservation do not disrupt mitochondrial function, they still have the ability to rescue the germ plasm, so both treatments can be used for subsequent experiments.

To explore the relationship between the quantity of mitochondria injected and the recovery of PGCs, we tested different numbers of injected mitochondria. Specifically, 2, 4, 8, 15 and 29 ×10^6^ mitochondria were injected into the vegetal pole of individual embryos. The number of PGCs increased progressively with concentrations from 2 to 8 ×10^6^ (Fig. 2D, right), reaching levels comparable to the control group. However, when 15 ×10^6^ mitochondria were injected, the number of PGCs began to decline, and at the highest concentration of 29 ×10^6^ mitochondria, embryos exhibited no detectable PGCs and experienced high rates of mortality.

In addition, to validate that only the injection of mitochondria-containing suspension from Percoll separation could rescue PGCs, we injected suspensions of the other bands (Supplementary Fig. 1) after eggs mitochondrial purification. As anticipated, none of these suspensions demonstrated the rescuing effect. These results therefore further support our conclusion that only mitochondria are capable of rescuing germ plasm.

### 3. Mitochondria labelled with PKH26 can be successfully tracked

The mitochondria were labelled with PKH26 and injected together with FITC-dextran into the vegetal pole of UV-irradiated fertilized embryos. PKH26-labelled mitochondria were colocalized in FITC-labelled PGCs at the tail-bud stage (Fig. 3A) suggesting that PGCs contained injected mitochondria. This was further confirmed by histology. In the sections, FITC-labelled PGCs and PKH26-labelled mitochondria were seen at the genital ridge of hatched larvae (Fig. 3B). Histological observation implies the successful *in vivo* tracking of labelled foreign mitochondria *via* PKH26 (Fig. 3C).

**Fig. 3.**
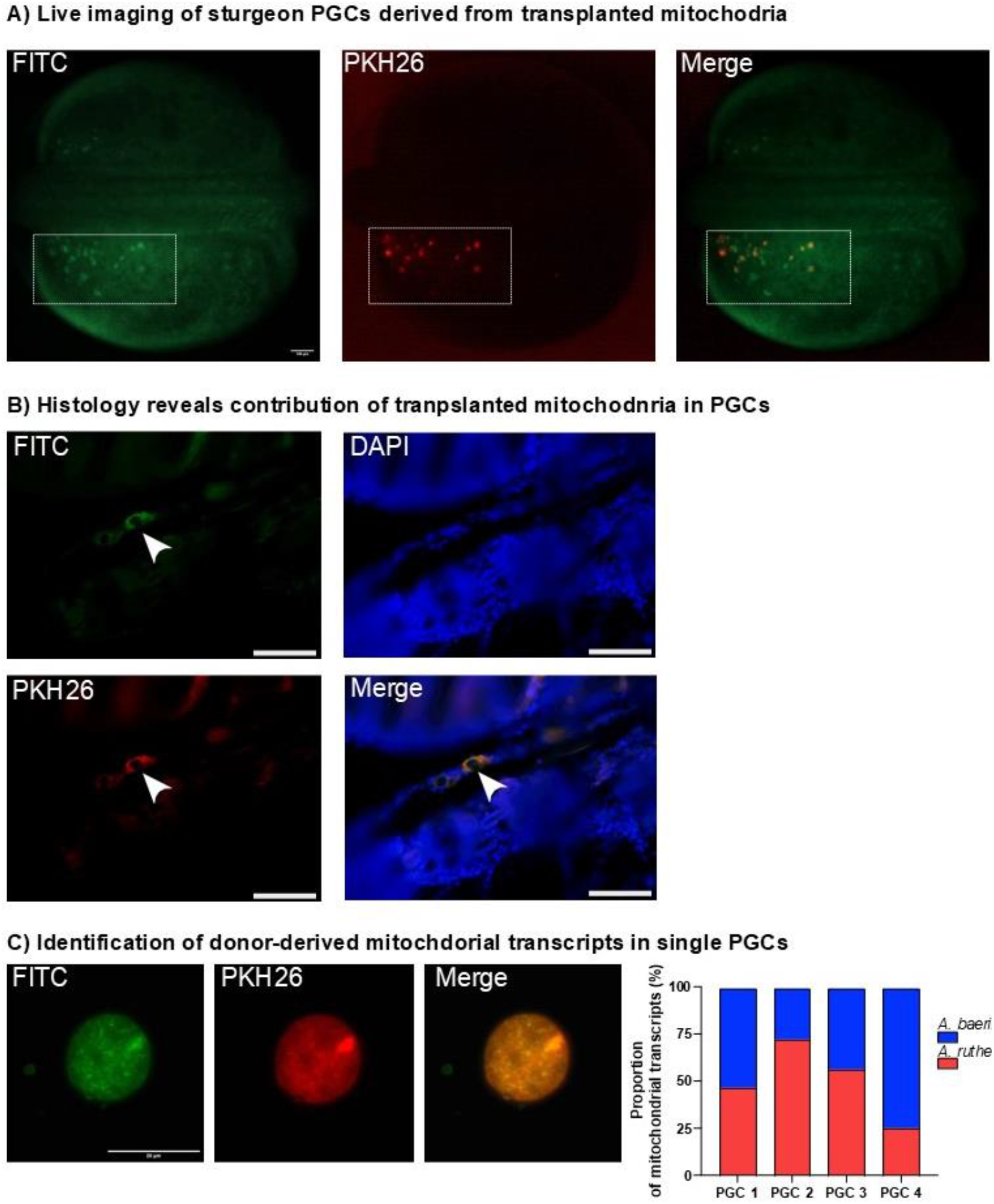
Interspecific transplanted mitochondria can be successfully tracked and identified. (A) At the tail-bud stage. FITC: FITC-dextran labelled PGCs. PKH 26: PKH26 labelled mitochondria formed PGCs. Merge: Merged image for FITC and PKH 26. The boxed areas show the PGCs. (B) In the genital ridge in larval at 12 days post fertilization (dpf). Merge: Merged image for FITC, PKH 26 and DAPI. The arrows show single PGC. (C) Left: Single PGC rescued with mitochondria injection. FITC: Single PGC stained with FITC. PKH 26: Single PGC formed by PKH26 labelled mitochondria. Merge: Merged image for FITC and PKH 26. Right: Proportion of mitochondrial transcripts in a single PCG. Scale bars in (A) indicate 100 μm, 10 μm in (B) and 20 μm in (C).

### 4. Identification of mitochondria after interspecific transplantation

The mitochondria isolated from Siberian sturgeon (*Acipenser baerii*) eggs were transplanted into sterlet (*Acipenser ruthenus*) embryos and identified with *A. ruthenus* mtDNA specific primers: 5’ - TATACACCATTATCTCTATGT - 3’ and 5’ - GGGAATAACCGTTAATTTGG - 3’, and *A. baerii* mtDNA specific primers: 5’ - TATACACCATTATCTCTATGT - 3’ and 5’ - CAGATGCCAGTAACAGGCTGA - 3’ (Mugue et al., 2008) at the larval stage. The mtDNA-specific PCR analysis showed positive amplification in all three analysed larvae (Supplementary Fig. 2).

To determine the proportion of endogenous and transplanted mitochondria in a single PCG, we performed ddPCR with mtDNA specific primers (*A. ruthenus* and *A. baerii*: 5’ - CACCATAATAACCGCCTTC - 3’ and 5’ - CRCCTCAGATTCATTGTAC - 3’) and probes (*A. ruthenus*, FAM: CGTCATCACCAATCTACTTTCCGC; *A. baerii*, HEX: CGTCATCACCAACCTCCTCTCCGC). The results are shown in Fig. 3C (right, details in Supplementary Figs 3 and 4). In single PGCs from sterlet larvae transplanted with Siberian sturgeon mitochondria, a high proportion of transplanted mitochondria were detected using probes specific to sterlet and Siberian sturgeon, accounting for 50.23% and 49.77% of mitochondrial transcripts, respectively. This clearly demonstrates that the mitochondria eliminated from germ plasm can be replaced by interspecific transplantation.

## Discussion

Our study highlights the critical role of mitochondria in germ plasm function and primordial germ cells (PGCs) specification in sturgeons. Using UV irradiation, we successfully depleted germ plasm, including mitochondria, in the vegetal pole of sturgeon embryos, leading to the complete loss of PGCs. Remarkably, transplantation of mitochondria isolated from donor eggs restored germ plasm function and rescued PGCs development. This demonstrates the indispensability of mitochondria in germline development. Importantly, we showed that mitochondria sourced from both the vegetal and animal poles of the egg, as well as purified mitochondria, restored germ plasm function to an equal extent, despite the absence of germ plasm determinants in the animal pole. These findings indicate that mitochondria in germ plasm are not uniquely specialized, but their presence is essential for PGCs development. These findings challenge prior assumptions that mitochondria within the germ plasm are uniquely specialized for germline determination and highlight their more generalizable functional role (Kloc et al., 2004).

The observation that mitochondria from eggs—but not from somatic tissues such as liver or muscle— successfully restored PGCs suggests that mitochondria from oocytes possess specific qualities, such as enhanced energy production and antioxidative capacity, that are critical for early embryonic development. Similar results have been observed in assisted reproduction, where mitochondria from oocytes are used to supplement defective mitochondria in species like humans, mice, and cattle (Friedman & Nunnari, 2014; Shoubridge & Wai, 2007; Van Blerkom, 2009, 2011). However, excessive mitochondrial transplantation in our study led to embryo sterility or death, indicating a physiological limit to mitochondrial supplementation. These findings underscore the necessity of precise control over mitochondrial transplantation procedures.

Studies in *Drosophila* (Ding et al., 1993) have demonstrated that certain mitochondrially encoded ribosomal RNAs function on the polar granules to specify the germ line. Similar observations have been made in *Xenopus* and the sea urchin, where mitochondrial ribosomal RNA has been detected in the germ plasm (Kashikawa et al., 2001; Kobayashi et al., 1998; Ogawa et al., 1999). Ikenishi and Kobayashi et al. (Ikenishi, 1998; Kobayashi et al., 1998) have further proposed that mitochondrial ribosomes within germ plasm may synthesize essential proteins necessary for the determination of germ line cells. However, mitochondrial ribosomes alone cannot achieve germ line cells’ differentiation, and additional determinants are required (Ephrussi et al., 1991; Kobayashi et al., 1998; Kobayashi & Okada, 1989; Satoru Kobayashi et al., 1993). For example, proteins such as Vasa, Tudor, and Piwi have been shown to regulate RNA localization and translation, while mRNA molecules like nanos and oskar are critical for specifying germline cells (Boswell & Mahowaldt, 1985; Cox et al., 1998; Ephrussi et al., 1991; Ephrussi & Lehmann, 1992; Tomancak et al., 1998). Furthermore, RNA-binding proteins like Dead end (*Dnd1*) contribute to the stabilization and proper expression of germline mRNAs (Weidinger et al., 2003).

It is noteworthy that previous studies on mitochondria and germ plasm have primarily focused on the relationship between mitochondria within the germ plasm and its associated components, with little attention given to the potential roles of mitochondria outside the germ plasm. Our experiments further underscore the critical contribution of mitochondria to germ plasm restoration and, for the first time, demonstrate that mitochondrial functionality is not dependent on their origin from the germ plasm. This finding highlights the broader functional importance of mitochondria and provides a novel platform for exploring unresolved questions regarding their contributions to germline specification.

The unique characteristics of the sturgeon germ plasm make sturgeon an ideal model organism for investigating the mechanisms by which mitochondria interact with germ plasm components to facilitate germline specification. Unlike teleost fish, where germ plasm determinants are distributed throughout the embryo during early development, sturgeons exhibit holoblastic cleavage, which spatially isolates the germ plasm from somatic lineages (Shah et al., 2022, 2024; Naraine et al., 2022). This isolation enabled targeted manipulations of germ plasm and its mitochondrial components without confounding effects from other embryonic tissues, creating ideal conditions for deeply uncovering the mechanisms by which mitochondria contribute to germline formation.

We employed multiple approaches to confirm the contribution of transplanted mitochondria to PGCs restoration. First, we quantified PGCs, showing that transplantation of donor mitochondria rescued PGCs numbers to control levels following UV irradiation. Second, we used PKH26 fluorescent cell tracing to successfully label transplanted mitochondria, enabling long-term tracking in host embryos. To our knowledge, this represents the first successful use of PKH26 for mitochondrial tracing in PGCs, exceeding tracking durations achieved with advanced methods like TPAP-C5-yne luminogens (Park et al., 2022). Third, we identified transplanted mitochondria using species-specific mitochondrial DNA (mtDNA) polymorphisms as described by Mugue et al. (Mugue et al., 2008). Finally, we used species-specific mitochondrial RNA (mtRNA) primers and digital PCR (dPCR) to confirm the presence of transplanted mitochondria in single PGCs. The use of mtRNA provided higher sensitivity and precision due to the greater abundance of RNA copies compared to DNA (Chen et al., 2023; Ozawa et al., 2007).

Our findings establish that mitochondria are indispensable for PGCs specification but do not require origin-specific properties from the germ plasm. This offers a new perspective on mitochondrial roles in germline biology and opens avenues for exploring mitochondrial-nuclear interactions. These results also hold potential applications in conservation genetics, particularly for critically endangered sturgeons. The transplantation of mitochondria into germ cells could complement other conservation techniques, such as androgenesis and nuclear transfer, to recover both maternal and paternal genetic information. Our laboratory has successfully demonstrated androgenesis in common carp (Cyprinus carpio) via cold shock (Kašpar et al., 2022), and efforts are underway to optimize this technique in sturgeon. Additionally, we have pioneered somatic cell nuclear transfer in sturgeon (Fatira et al., 2018, 2019), providing a foundation for advanced reproductive biotechnologies.

In parallel, alternative approaches to addressing the challenges of fish egg and embryo cryopreservation have been explored, focusing on germline stem cells. These include embryonic PGCs (Saito et al., 2008) or adult gonial stem cells (Yoshizaki & Lee, 2018). These specialized cells can be isolated from a target species and stored in liquid nitrogen. When required, the cells can be thawed and transplanted into suitable surrogate parents with depleted germ cells. Subsequently, donor-derived progeny is produced. This reproductive biotechnology is at present available for several fish species (de Siqueira-Silva et al., 2021; Franěk et al., 2021; Hattori et al., 2019; Kawamura et al., 2024; Okutsu et al., 2007; Yutaka Takeuchi et al., 2004). In sturgeon the transplantation of their gonial stem cells into interspecies recipients has been optimized (PšeniČka et al., 2015; Romney et al., 2023). Additionally, several methods for eliminating PGCs have been developed, including the use of antisense morpholino oligonucleotides and CRISPR/Cas9 gene editing targeting the *dnd*1 gene (Baloch et al., 2019; Linhartová et al., 2015), and UV irradiation targeting the germ plasm of embryos (Saito et al., 2018). Despite these significant advancements, the attainment of surrogate reproduction in sturgeons remains an unachieved goal due to the lack of an effective sterilization method to produce a large number of sterile hosts and the slow maturation of sturgeons.

In conclusion, this study provides compelling evidence of mitochondria’s essential role in germ plasm function and PGCs development. Our results contribute to a deeper understanding of germline biology and lay the groundwork for future research into mitochondrial-nuclear interactions, with potential applications in both evolutionary biology and conservation science.

## Supplementary Information

Document S1. Supplementary Figures 1 - 4 and Supplementary Note.

## Declaration of Interest

The authors declare that there is no conflict of interest that could be perceived as prejudicing the impartiality of the research reported.

## Funding

The work was supported by the Ministry of Education, Youth and Sports of the Czech Republic - project Biodiversity (CZ.02.1.01/0.0/0.0/16_025/0007370), and the Czech Science Foundation (grant number 22-31141J). This project has received funding from the European Union’s Horizon 2020 Research and Innovation Programme under grant agreement No. 871108 (AQUAEXCEL3.0). This study was also supported by 86652036 from RVO. This output reflects only the view of the author, and the European Union cannot be held responsible for any use that may be made of the information contained therein.

## Author contributions

M.P. conceptualized and supervised the study. R.S. designed the primers and probes for ddPCR and performed the ddPCR. T.T. designed primers for Sanger sequencing and analyzed Sanger products to identify pair-wise SNPs. R.F. edited the figures and manuscript. T.S. reviewed and edited the manuscript. L.G. performed the experiments, analysed the data, and wrote the draft manuscript. M.R. and D.G. helped with fish reproduction. All co-authors revised the manuscript and agreed to the final version. All authors have read and agreed to the published version of the manuscript.

## Acknowledgements

We sincerely thank John Craig for his assistant in correcting the English of this manuscript. We are deeply grateful to Shah Mujahid Ali for his assistance with embryo injections and histology work. We thank Eva Rohlova from the GeneCore facility in BIOCEV for assistance during ddPCR optimization and analysis. We are also thanking VazaČová Michaela, Linhartová Zuzana, Nayak Rigolin and the members of the Laboratory of Germ Cells, Faculty of Fisheries and Protection of Waters, University of South Bohemia in Ceske Budejovice, for their support on experiments. We acknowledge the BC CAS core facility LEM supported by MEYS CR (LM2023050 Czech-BioImaging and OP VVV CZ.02.1.01/0.0/0.0/18_046/0016045).

## Notes

### Competing Interest Statement

The authors have declared no competing interest.

