## Supplementary Figures 1-4 and Supplementary Note for "Mitochondria as Indispensable Yet Replaceable Components of Germ Plasm: Insights into Primordial Germ Cell Specification in Non-Teleost Sturgeons"

**Inventory of Supplementary Information**

**Supplementary Figures**

**Figure 1:** Crude and purified mitochondria.

**Figure 2:** Species specific PCR confirms presence of donor mtDNA after PGCs rescue.

**Figure 3:** Validation of reaction specificity of mtDNA specific primers and probes for Siberian sturgeon and sterlet.

**Figure 4:** Concentration in single PGC used for identification from sterlet larvae transplanted with Siberian sturgeon mitochondria.

**Supplementary Note**

**Supplementary Figures**

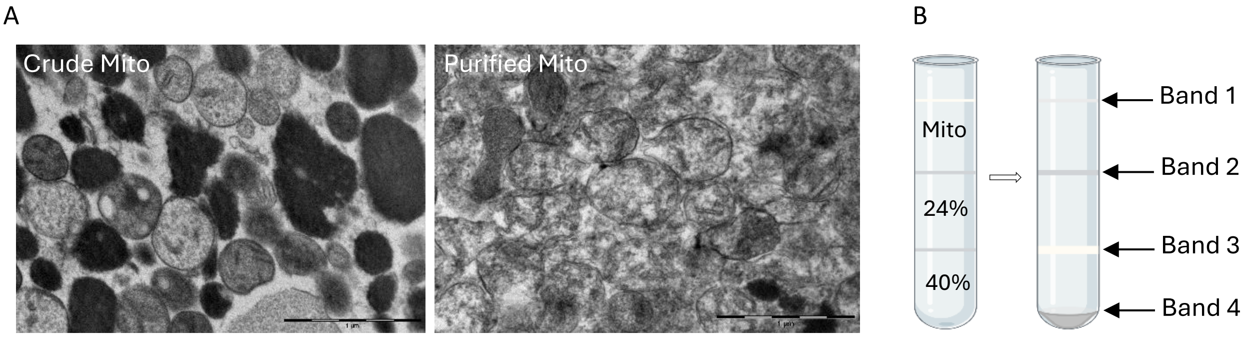


**Figure 1. Crude and purified mitochondria.** (A) TEM images. (B) Percoll density gradients for the purification of mitochondria. Preparations of mitochondria on a step gradient consisting of an upper 24 % and a lower 40 % Percoll layer (left). 4 bands were observed after centrifugation of crude mitochondria (right). The purified mitochondria were collected from band 3 (between 24 and 40 % Percoll).

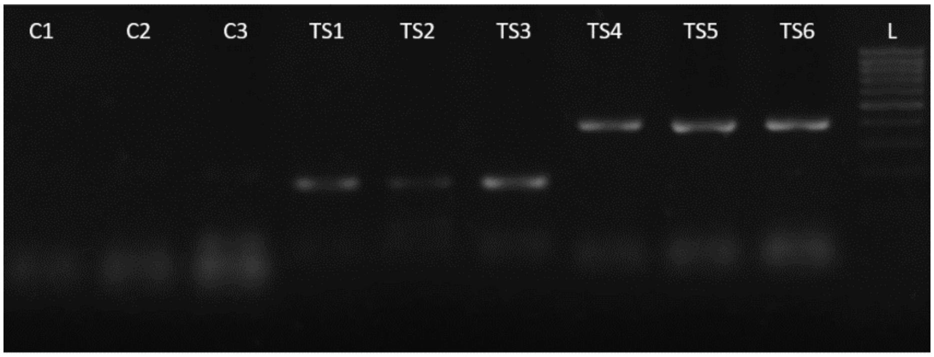


**Figure 2. Species specific PCR confirms presence of donor mtDNA after PGCs rescue.** C1-3: Sterlet larvae using Siberian sturgeon mtDNA specific primers show no PCR amplification. TS1-6: Sterlet embryos after UV irradiation transplanted by Siberian sturgeon mitochondria. TS1-3: Using sterlet mtDNA specific primers, it shows positive amplification. TS4-6: Using Siberian sturgeon mtDNA specific primers, it shows positive amplification. L: 1000bp DNA marker.

A B

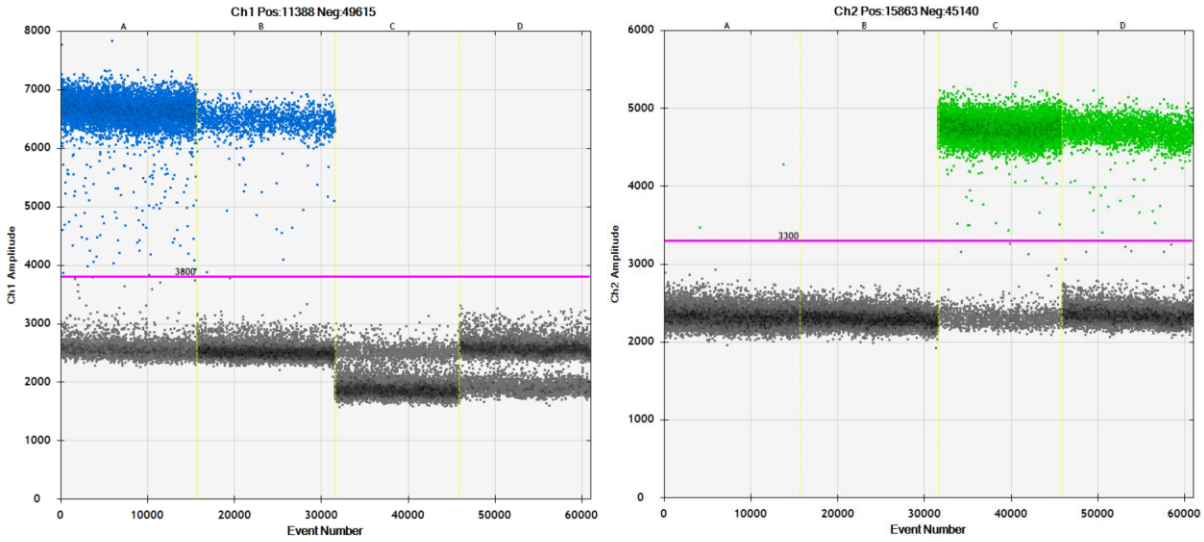


**Figure 3. Amplitude images for validating reaction specificity.** Samples A through D are 50 pg of Sterlet DNA, 5 pg of Sterlet DNA, 50 pg of Siberian DNA and 5 pg of Siberian DNA, respectively. (A) FAM channel with sterlet primers and probes, it shows positive amplification in sterlet samples. (B) HEX channel with Siberian sturgeon primers and probes, it shows positive amplification in Sibrian samples.

A B

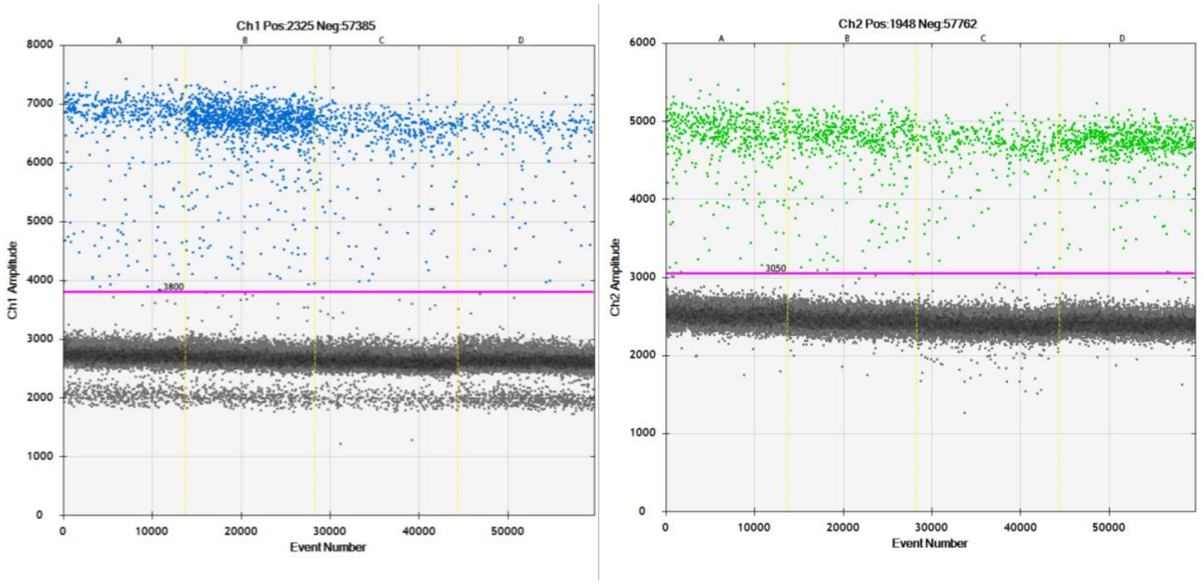


**Figure 4. Amplitude images of concentrations in single PGC.** Samples A through D are single PGC isolated from transplanted sterlet larvae with Siberian sturgeon mitochondria. (A) FAM channel with sterlet primers and probes, it shows positive amplification. (B) HEX channel with Siberian sturgeon primers and probes, it shows positive amplification. dPCR mixture contained DNA from lysed single PGC collected in 5 µl of RNase-free distilled water (ThermoFisher, USA) and the remaining steps were performed as in validation.

**Supplementary Note**

**ddPCR primer and probe design sequence (Sterlet different from Siberian is highlighted with yellow, Siberian different from Sterlet is highlighted with turquoise).**

TTGTCACACAAATCCTAACAGGACTATTTCTCGCAATACACTACACAGCTGACATTTCAWCAGMCTTCTCCTCTGTCGCCCA
CATCTGCCGAGATGTAAATTACGGATGATTAATCCGCAATATTCATGCAAACGGGGCCTCCTTCTTCTTTATCTGCTTGTATCT
CCACGTAGCACGAGGCATGTACTACGGTTCTTATCTCCAAAAAGAAACCTGAAACATCGGAGTAATCCTCCTGCTCCTCACC
ATAATAACCGCCTTCGTAGGATATGTACTACCCTGAGGACAGATATCATTTTGAGGAGCAACCGTCATCACCAATCTACTTTC
CGCCTTTCCGTACATCGGCGACACACTAGTACAATGAATCTGAGGCGGCTTTTCAGTAGACAACGCCACCCTCACCCGATT
TTTCGCCTTTCACTTTCTTCTACCATTCGTAATCGCCGGAGCTAGCATAATTCACCTTCTGTTTCTACACCAAACAGGATCA
